# A metabolic dependency for host isoprenoids in the obligate intracellular pathogen *Rickettsia parkeri* underlies a sensitivity for the statin class of host-targeted therapeutics

**DOI:** 10.1101/528018

**Authors:** Vida Ahyong, Charles A. Berdan, Daniel K. Nomura, Matthew D. Welch

## Abstract

Gram-negative bacteria in the order Rickettsiales are obligate intracellular parasites that cause human diseases such typhus and spotted fever. They have evolved a dependence on essential nutrients and metabolites from the host cell as a consequence of extensive genome streamlining. However, it remains largely unknown which nutrients they require and whether their metabolic dependency can be exploited therapeutically. Here, we describe a genetic rewiring of bacterial isoprenoid biosynthetic pathways in the Rickettsiales that has resulted from reductive genome evolution. We further investigated whether the spotted fever group *Rickettsia* species *Rickettsia parkeri* scavenges isoprenoid precursors directly from the host. Using targeted mass spectrometry in uninfected and infected cells, we found decreases in host isoprenoid products and concomitant increases in bacterial isoprenoid metabolites. Additionally, we report that bacterial growth is prohibited by inhibition of the host isoprenoid pathway with the statins class of drugs. We show that growth inhibition correlates with changes in bacterial size and shape that mimic those caused by antibiotics that inhibit peptidoglycan biosynthesis, suggesting statins inhibit cell wall synthesis. Altogether, our results describe an Achilles’ heel of obligate intracellular pathogens that can be exploited with host-targeted therapeutics that interfere with metabolic pathways required for bacterial growth.

**Importance:** Obligate intracellular parasites, which include viruses as well as certain bacteria and eukaryotes, extract essential nutrients and metabolites from their host cell. As a result, these pathogens have often lost essential biosynthetic pathways and are metabolically dependent on the host. In this study, we describe a metabolic dependency of the bacterial pathogen *Rickettsia parkeri* on host isoprenoid molecules that are used in the biosynthesis of downstream products including cholesterol, steroid hormones, and heme. Bacteria make products from isoprenoids such as an essential lipid carrier for making the bacterial cell wall. We show that bacterial metabolic dependency can represent an Achilles’ heel, and that inhibiting host isoprenoid biosynthesis with the FDA-approved statin class of drugs inhibits bacterial growth by interfering with the integrity of the cell wall. This work highlights a potential to treat infections by obligate intracellular pathogens through inhibition of host biosynthetic pathways that are susceptible to parasitism.

## Introduction

Gram-negative alphaproteobacteria in the family Rickettsiaceae, which includes the genera *Rickettsia* and *Orientia*, are obligate intracellular parasites that can cause human diseases including spotted fever (spotted fever group (SFG) *Rickettsia*), typhus (typhus group (TG) *Rickettsia*), and scrub typhus (*Orientia* species) (1). These bacteria are transmitted to mammals by arthropod vectors such as fleas, ticks, and mites. Although most pathogenic species cause moderately severe illnesses, in some cases infections can be fatal if untreated or even upon delayed treatment with first-line antibiotics (2). We study the SFG species *R. parkeri,* which causes an eschar at the site of the tick bite and symptoms that include fever, malaise, nausea, headaches, and an occasional rash (3, 4).

Upon invasion of host cells, the Rickettsiaceae quickly escape the primary vacuole into the host cell cytoplasm, where they grow and proliferate. One property of obligate intracellular bacteria such as *Rickettsia* is that adaptation to a dependency upon growth inside host cells has resulted in genome size reduction (5-9). The relatively small genomes of *Rickettsia* species (∼1.1-1.5 Mbp) encode for a reduced number of proteins (1273 predicted proteins in *Rickettsia parkeri;* NCBI reference sequence: NC_017044.1). This typically correlates with the loss of genes encoding components of metabolic biosynthetic pathways, together with the requirement to scavenge essential metabolites from the host (5, 8).

One essential class of metabolites are the isoprenoids (also known as terpenoids), which are derived from simple five carbon isoprene units that are assembled to make thousands of different molecules. The biosynthesis of the central isoprene precursor molecule, isopentenyl pyrophosphate (IPP), and its isomer, dimethylallyl diphosphate (DMAPP), occurs through two distinct pathways. The mevalonate (MEV) pathway (predominantly in archaea, Gram-positive bacteria, and animals) or the 2-C-methyl-D-erythritol 4-phosphate (MEP) pathway (primarily in Gram-negative bacteria) (Figure 1). In bacteria, isoprenoid biosynthesis produces vital products such as bactoprenols (essential building blocks for peptidoglycan (PG) and other cell wall polysaccharides), ubiquinone or menaquinone (involved in the electron transport chain), or chlorophylls in cyanobacteria (10). In mammalian cells, isoprenoid biosynthesis produces a larger variety of products including cholesterol, ubiquinone, steroid hormones, prenylated proteins, heme, and vitamin K (11).

**Figure 1:**
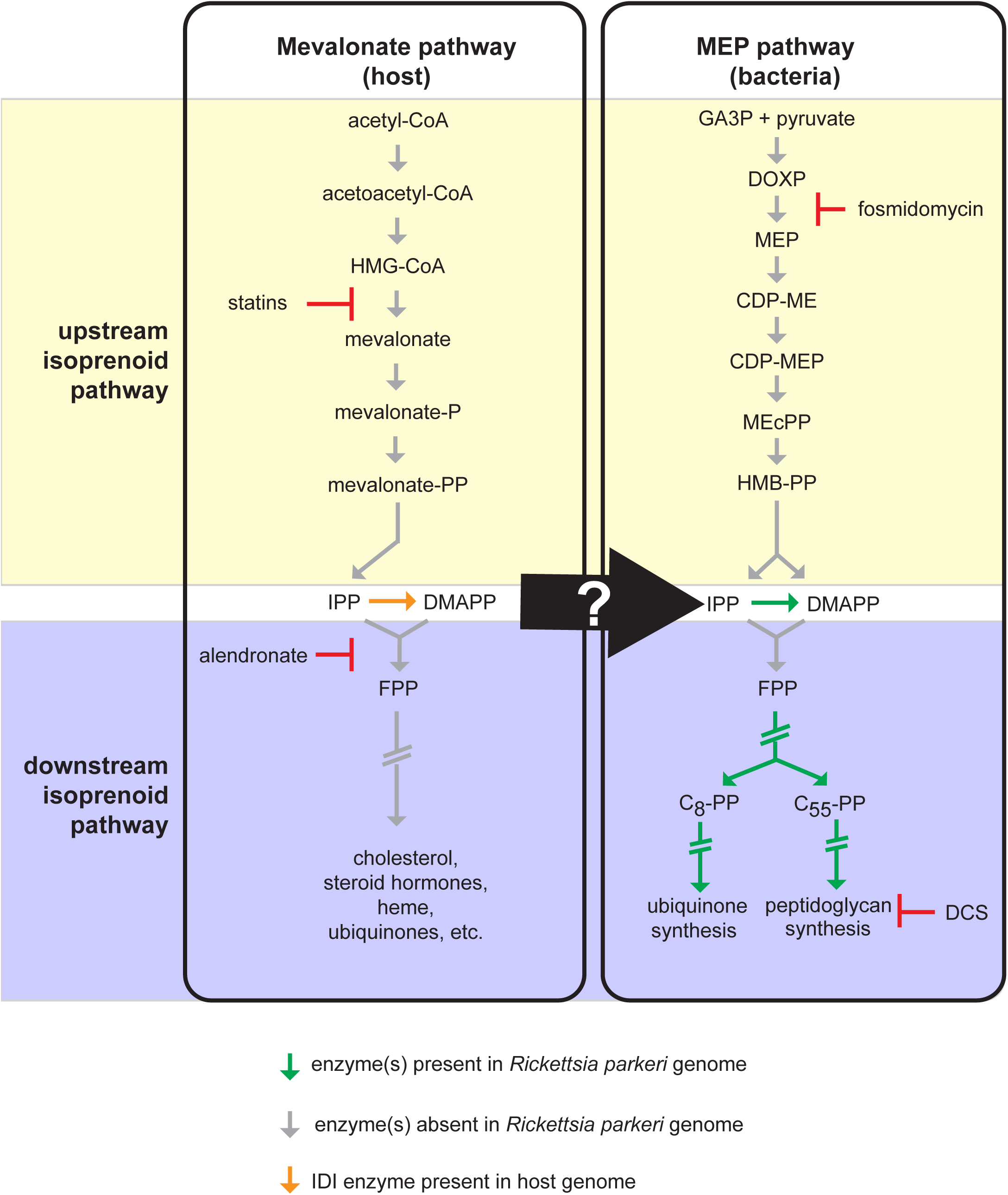
Generalized MEV and MEP pathways leading to downstream isoprenoid products. The green arrows indicate the presence of annotated enzymes encoded in the *R. parkeri* genome. The gray arrows indicate the absence of annotated enzymes encoded in the *R. parkeri* genome. The orange arrow indicates the presence of the IDI enzyme from the host genome. The light yellow background denotes the upstream isoprenoid pathway whereas the light blue denotes the downstream isoprenoid pathway. The central black arrow and white question mark denote a possible transporter for isoprenoids from the host into the bacteria. Double hash arrows indicate multiple enzymatic steps. Statin drugs inhibit the activity of HMG-CoA reductase and the formation of mevalonate of the MEV pathway. Fosmidomycin is an inhibitor of DXP reductoisomerase of the MEP pathway. Alendronate sodium hydrate is an inhibitor of farnesyl diphosphate synthase. D-cycloserine (DCS) is an inhibitor of D-alanine racemase and D-alanine-D-alanine ligase.

Although the isoprenoid biosynthetic pathway is essential, its components are missing from certain obligate pathogens including *Mycoplasma* species (12) and the protozoan parasite *Cryptosporidium* (13). Bioinformatic analysis (14) shows that genes encoding the upstream enzymes from the MEV and MEP pathways that are required to make IPP and DMAPP are also missing in *Rickettsia* species, with the exception of the *idi* gene encoding the enzyme isopentenyl diphosphate isomerase (IDI), which catalyzes the reversible conversion of IPP to DMAPP (15). Since *Rickettsia* have access to metabolites in the host cytoplasm, this suggests that *Rickettsia* may steal IPP and/or DMAPP, altering the flux of host-derived isoprenoid precursors to initiate downstream bacterial isoprenoid biogenesis. The downstream enzymes required to utilize these short-chained isoprenoid precursors for ubiquinone biosynthesis and PG biosynthesis are encoded in *Rickettsia* genomes.

In this study, we explore the evolutionary changes that occurred within the order *Rickettsiales* with respect to the presence of genes encoding components of the isoprenoid biosynthetic pathway. We also show that bacterial infection results in depletion of host isoprenoid products and the synthesis of bacterial isoprenoid products. Furthermore, we use a chemical genetic approach to reveal that inhibition of host isoprenoid biosynthesis by statin treatment prevents bacterial growth and alters bacterial shape. Our results suggest that the host MEV pathway serves as the upstream source of isoprene units for the synthesis of bacterial bactoprenols and ubiquinone, and that inhibition of the host MEV pathway by statins may represent a promising avenue for host-directed therapeutics to treat *Rickettsia* infection.

## Results

### The isoprenoid biosynthesis pathway is under evolutionary flux in the order Rickettsiales

We first sought to determine the evolutionary conservation of the isoprenoid biosynthesis pathway in the order *Rickettsiales*. We constructed a phylogenetic tree using 16S rDNA sequences across six main genera of the order as well as the closest free-living organism, *Pelagibacter ubique,* and assessed the presence or absence of genes encoding upstream or downstream components of the MEP pathway as well as the IDI enzyme (Figure 2A). We confirmed that the *Rickettsia* (14) and *Orientia* lineages lack the genes encoding upstream MEP pathway enzymes but have those encoding the downstream pathway components, whereas the other genera contained the full complement of MEP pathway genes. None have MEV pathway genes, with the exception of the *idi* gene in *Rickettsia* species. The absence of upstream MEP pathway genes suggests that *Rickettsia* must scavenge MEV/MEP pathway intermediates (IPP, DMAPP or FPP, Figure 1) from the host cell.

**Figure 2:**
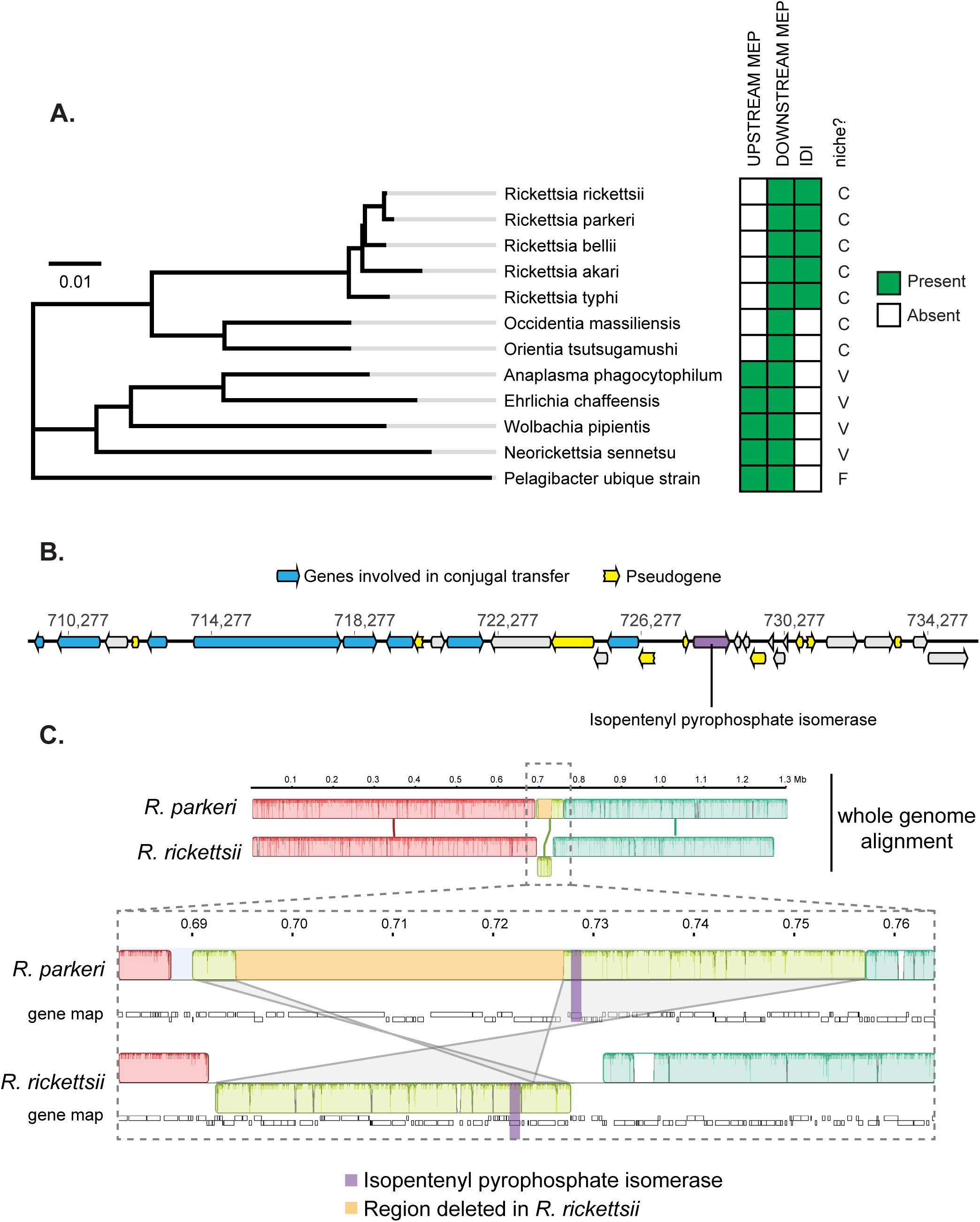
Genome evolution in the *Rickettsiales* isoprenoid pathway. **A.** Phylogenetic neighbor-joining tree of the *Rickettsiales* and the outgroup *Pelagibacter ubique.* Scale bar length represents 0.01 nucleotide substitutions per site. The presence of intact upstream MEP pathway genes or *idi* gene for each species is indicated by a green-filled box. The environmental niche is indicated in the adjoining column, as follows: C, intracellular cytoplasmic; V, intracellular vacuolar; F, free living. **B.** A schematic of the *R. parkeri idi* genome locus. Genes involved in conjugal transfer are colored in blue. Genes that are pseudogenized are in yellow. The *Idi* gene is highlighted in purple. **C.** Whole genome alignment of *R. parkeri* and *R. rickettsii*. The *R. rickettsii* genome has a large 33.5kb deletion (in orange) in the genome alignment for the region adjacent to the *idi*. The gray line segments indicate inverted alignment orientation between *R. parkeri* and *R. rickettsii.*

Additionally, the *idi* gene is only present in the *Rickettsia* lineage but not the *Orientia* lineage. A previous report suggested that the *idi* gene in *Rickettsia* was acquired by horizontal gene transfer (14). To further investigate this, we examined the genomic locus surrounding the *idi* gene in the *R. parkeri* str. Portsmouth genome (Figure 2B). We found many surrounding genes that were associated with conjugal transfer functions, such as transposases and pili. Furthermore, there are nine surrounding pseudogenes with premature stop codons, suggesting this region is in flux and may be in the process of undergoing genomic reduction. To further test this hypothesis, we performed a whole genome alignment between *R. parkeri* and the closely related yet more pathogenic species *R. rickettsii*. Interestingly, we found the two genomes were highly syntenic, with the exception of a 65kb region containing the *idi* gene (Figure 2C, region in lime green, *idi* gene in purple) that is inverted between the two species and in the *R. rickettsii* genome contains a 33.5 kb deletion (orange highlighted region in Figure 2C and Supplemental Table S1). These observations suggest that the locus surrounding the *idi* gene was acquired by horizontal gene transfer and is still under evolutionary pressure to retain the *idi* gene while simultaneously undergoing continuing reduction.

### *R. parkeri* infection results in depletion of host isoprenoid products and accumulation of bacterial isoprenoid products

To test whether *R. parkeri* scavenges host isoprenoids to make bacterial products, we measured the presence and abundance of host and bacterial isoprenoid-derived metabolites by liquid chromatography-mass spectrometry (LC-MS/MS). Confluent infected and mock-infected cultures of African green monkey kidney epithelial Vero cell cultures were incubated for 4 d, collected and then extracted for lipid metabolites. For host isoprenoids, we monitored cholesterol, total cholesteryl esters, cholesteryl oleate, and ubiquinone-10 using single-reaction monitoring (SRM)-based LC-MS/MS methods (Figure 3). We observed a statistically significant approximately 2-fold decrease in total cholesteryl esters and cholesteryl oleate in the infected compared with uninfected samples. These products are generally understood to be the storage units of cholesterol packaged in intracellular lipid droplets (16). In contrast, we saw no change in cholesterol or ubiquinone-10. Thus, infection results in the depletion of isoprenoid-derived storage forms of cholesterol.

**Figure 3:**
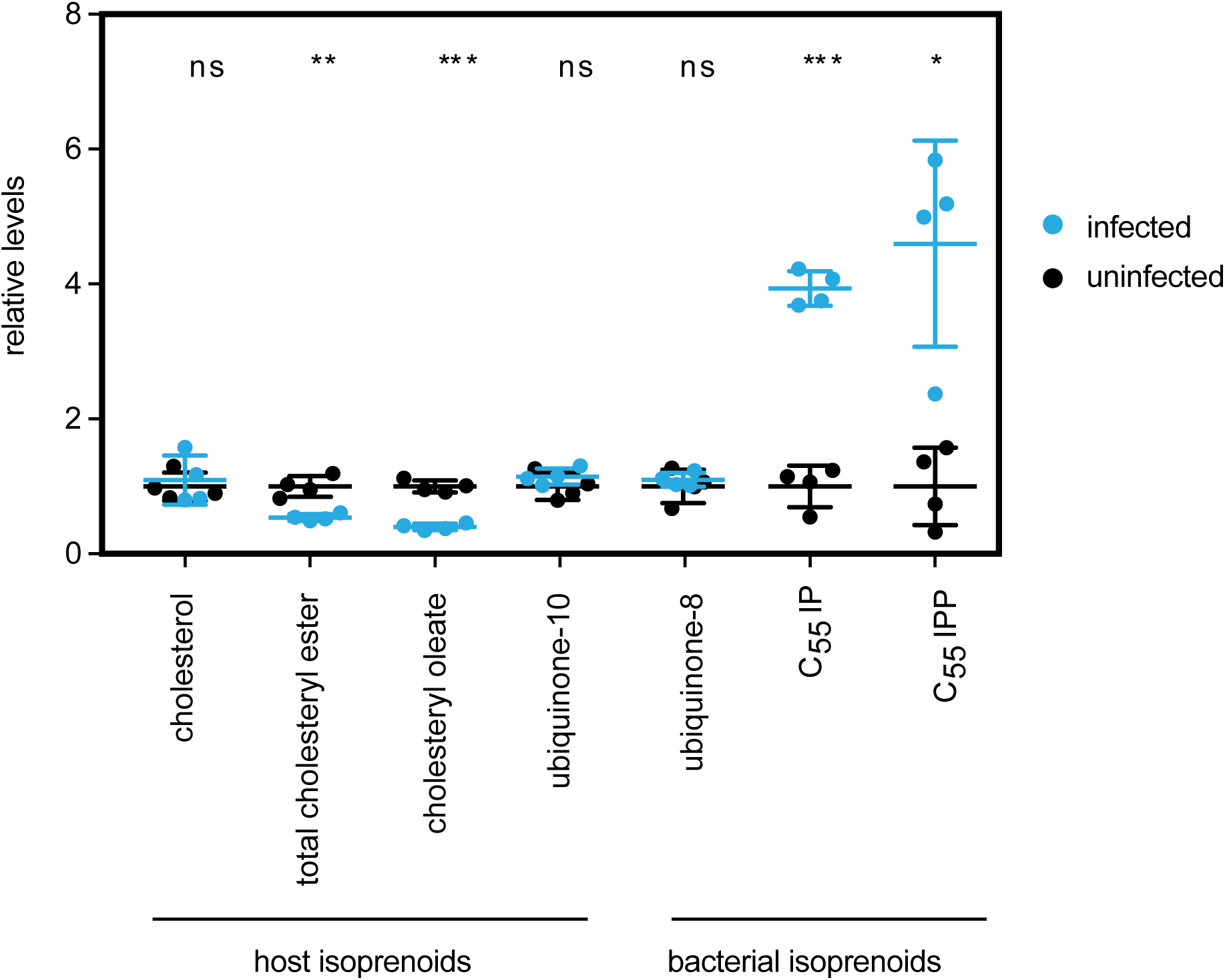
Targeted mass spectrometry of bacterial and host isoprenoids. Graph of the relative levels of host and bacterial isoprenoids in 4 d.p.i. uninfected and infected Vero cell cultures, normalized to internal standards. Four replicates were done for each experiment. Error bars represent standard deviation. Statistical comparisons were done by the unpaired Student’s t-test; ns, not significant; *, p<0.05; **, p<0.01; ***, p<0.001.

For bacterial isoprenoids, we measured bactoprenols, which include the isoprenoid products C_55_-isopentyl pyrophosphate (C_55_-IPP) and C_55_-isopentyl phosphate (C_55_-IP). C_55_-IPP is initially produced by a dedicated prenyltransferase, UppS, and must be dephosphorylated to C_55_-IP to act as a lipid carrier (17). We also measured bacterial ubiquinone, ubiquinone-8, which contains an 8 prenyl subunit tail, in contrast with the human version containing 10 prenyl subunits, ubiquinone-10 (Figure 3) (18, 19). We observed a statistically-significant approximately 4-fold increase in both C_55_-IPP and C_55_-IP in infected compared with uninfected cells. We did not, however, find significant differences in bacterial ubiquinone (ubiquinone-8), possibly due to the presence of ubiquinone-8 intermediates from the host cell ubiquinone-10 biosynthesis pathway.

These results indicate that bacterial isoprenoid products, primarily bactoprenols, accumulate during *R. parkeri* infection. Collectively, the depletion of host isoprenoid products and accumulation of bacterial isoprenoid products, even in the absence of a bacterial isoprenoid synthesis pathway, suggests that isoprenoids are scavenged from the host by *R. parkeri*.

### Chemical inhibition of the host mevalonate pathway inhibits bacterial growth

To determine whether host isoprenoids are necessary for bacterial growth, we sought to inhibit the host mevalonate pathway and reduce the pool of available host IPP and DMAPP. We used statins, a class of drugs that block the activity of HMG-CoA reductase (Figure 1). A previous report showed that pre-incubation of *R. conorii* infected mouse L929 fibroblast cells with the statin lovastatin caused a reduction in *R. conorii* plaque size, which was interpreted to result from a reduction in cholesterol-dependent adherence of bacteria to host cells (20). To bypass possible effects of statins on bacterial adherence and invasion, we allowed infection of Vero cells to proceed for 2 h to allow for maximal bacterial adherence/invasion (21), treated cells with (or without) various concentrations of the statin pitavastatin, and then measured bacterial numbers using a *R. parkeri-*specific qPCR endpoint assay (Figure 4A). We observed a >2-log reduction in bacterial growth with increasing pitavastatin concentrations. To ensure that statin inhibition of bacterial growth was dependent on HMG-CoA reductase inhibition by pitavastatin and not secondary effects of the drug, we tested whether the downstream product of HMG-CoA reductase, mevalonate, could rescue this growth inhibition. Indeed, co-treated with 400 µM mevalonate along with pitavastatin rescued bacterial growth (Figure 4A).

**Figure 4:**
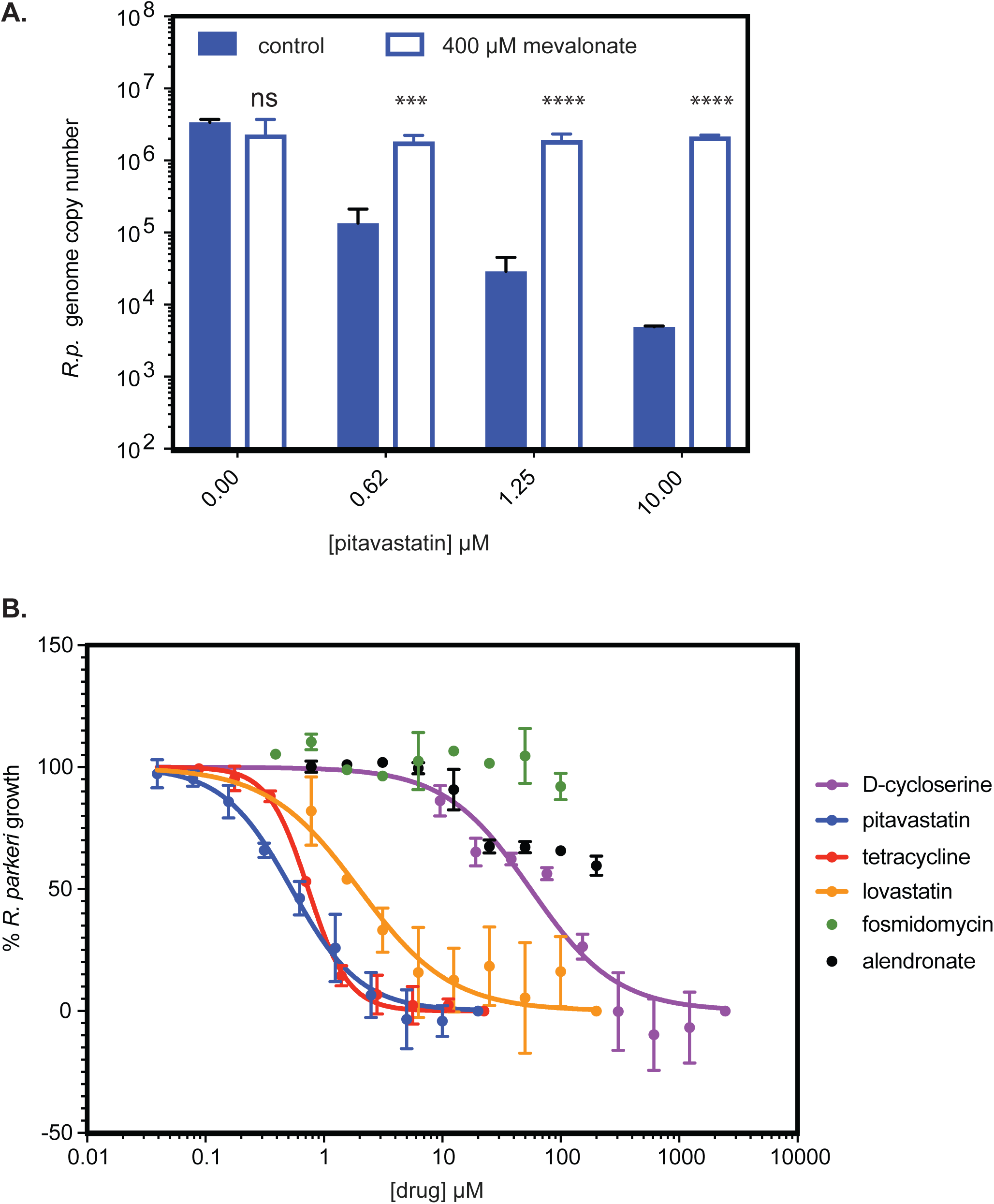
Chemical inhibition and rescue of *R. parkeri* growth. **A.** Graph of *R. parkeri* genome copy numbers in the presence of varying concentrations of pitavastatin, without or with mevalonate. Four replicates were done, and error bars represent standard deviation. Statistical comparisons were done by the unpaired Student’s t-test for each concentration of pitavastatin for the wild type vs wild type with mevalonate supplementation; ns, not significant; *, p<0.05; **, p<0.01; ***, p<0.001; ***, p<0.0001. **B.** Dose-dependent growth inhibition of *R. parkeri* in the presence of the indicated drugs targeting MEV or MEP pathway enzymes, or tetracycline, normalized to a no-drug control. Fosmidomycin and alendronate failed to generate fit curves. Measurements for each drug concentration were performed in duplicate and error bars represent standard deviation.

To further assess the importance of the MEV pathway, we generated dose-response curves for the effect of lovastatin and pitavastatin, which are chemically distinct, on *R. parkeri* growth using a 96-well qPCR endpoint assay. We observed a dose-dependent inhibition of bacterial growth with lovastatin (EC_50_ = 2.0 µM) and even more robust inhibition with pitavastatin (EC_50_ = 0.5 µM) (Figure 4B). This data indicates that the host MEV pathway is critical for *R. parkeri* growth.

To further test that bacterial growth is dependent on a complex integration of the host and pathogen isoprenoid pathways (as depicted in Figure 1), we tested for dose-dependent growth inhibition using additional inhibitors of MEV and MEP pathway enzymes (Figure 4B). Fosmidomycin, which specifically inhibits 1-deoxy-D-xylulose 5-phosphate (DXP) reductoisomerase, a key enzyme in the bacterial MEP pathway, caused no inhibition of bacterial growth at up to 100 µM. Alendronate sodium, a specific inhibitor of host farnesyl diphosphate (FPP) synthase, caused approximately 50% growth inhibition. This further suggests that *R. parkeri* growth is independent of bacterial isoprenoid production, and partially dependent on FPP from the host.

Finally, we tested the anti-bacterial antibiotics tetracycline, which is used as a first-line treatment for *Rickettsia* infection and targets protein synthesis, or D-cycloserine, which blocks the PG biosynthesis pathway that also requires isoprenoid biosynthesis products. Tetracycline inhibited growth with an EC_50_ = 0.7 µM, consistent with minimal inhibitory concentrations (MIC) values for SGF rickettsiae of 0.06-0.25 µg/mL (0.1-0.5 µM) (22). D-cycloserine also caused a dose-dependent inhibition of *R.parkeri* growth with an EC_50_ = 56 µM, consistent with effects on *R. prowazekii* (23), and with minimal inhibitory concentrations (MIC) values for *Escherichia coli* K12 (MIC = 30 µg/mL, 290 µM) (24) and *Mycobacterium tuberculosis* (15 µg/mL, 150 µM) (25). Taken together, the results of chemical inhibition assays support the notion that *Rickettsia* are dependent on the upstream host MEV pathway, but not the upstream MEP pathway. Furthermore, they are susceptible to inhibition of PG biosynthesis.

### Statins treatment causes bacterial shape defects that mimic those caused by peptidoglycan-targeting antibiotics

We sought to further assess the effect of statin treatments on bacterial physiology. Limiting the availability of isoprenoids is predicted to result in reduced PG due to the requirement for isoprenoids in PG synthesis. Though *Rickettsia* (22)*1*, like other Gram-negative bacteria (26), are generally resistant to PG targeted antibiotics, one study showed that high concentrations of penicillin produced spheroplasts (27), a type of L-form bacteria that have a defective PG cell wall and take on a spherical shape (28). Therefore, we sought to measure whether statin inhibition of the host MEV pathway altered bacterial shape in a similar manner to inhibition of PG synthesis. To this end, we performed immunofluorescence microscopy on 96-well plates of infected cells at 3 dpi to track the alterations of bacterial shape in response to dose-dependent application of statins, D-cycloserine, or tetracycline as a control. We performed automated image analysis using CellProfiler (29) and measured a number of bacterial cell shape features. We determined that two shape measurements, area and eccentricity (circle = 0, rod/line = 1), accounted for most of the shape alterations seen in the concentration ranges between EC_50_-EC_100_. In D-cycloserine treated cells, there was a dose-dependent increase in area and decrease in eccentricity as bacteria became more circular and less rod-shaped (Figure 5A, B). In cells treated with either lovastatin or pitavastatin, similar changes in area and eccentricity were seen along the dose-response curve. In contrast, upon treatment with tetracycline, we found little change in the bacterial area, although there was a general decrease in eccentricity.

**Figure 5:**
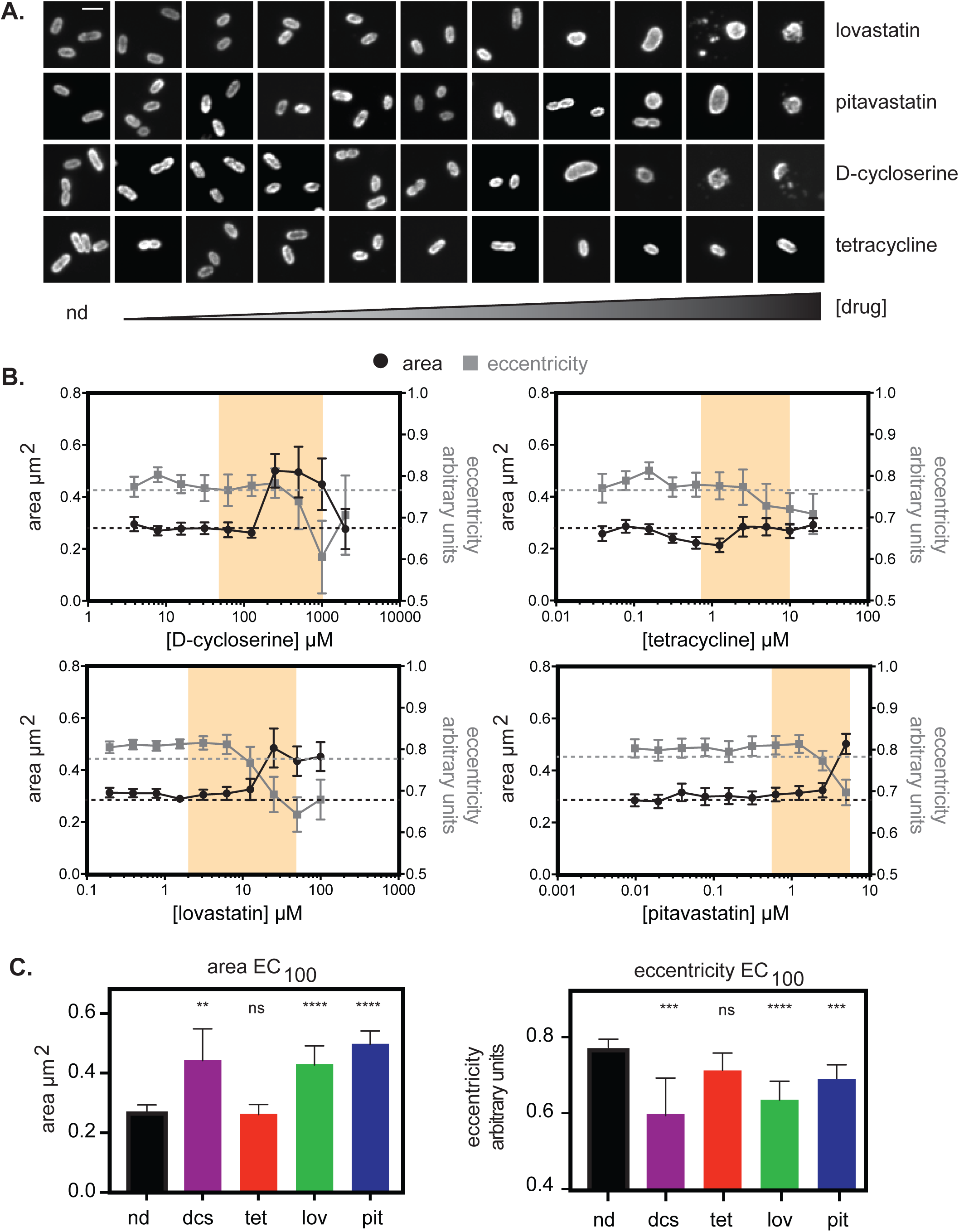
Shape and size measurements of *R. parkeri* bacteria under different drug pressures. **A.** Representative bacterial cells from the EC curve drug concentrations from Figure 5B. Scale bar = 1 µm. **B.** Dose-dependent changes in area and eccentricity of *R. parkeri* bacterial cells under drug treatment with lovastatin, pitavastatin, tetracycline, or D-cycloserine. Each measurement is the mean of at least 40 individual bacterial cells, except for concentrations of D-cycloserine >250 µM where most bacterial cells were lysed and fewer measurements were possible (for 500 µM DCS, n=22; for 1000 µM DCS, n=18; for 2000 µM DCS, n=13). Error bars represent the 95% confidence interval. **C.** Graphs plotting area and eccentricity measurements at the EC_100_ values calculated from Figure 4B and matched to measurements from Figure 5A. Regions shaded in yellow are the calculated EC_50_-EC_100_ values from Figure 4B. Graphs show the mean of the sample set and error bars represent the 95% confidence interval. Statistical comparisons were done by one-way ANOVA of all drug treated samples compared to the no drug (nd) sample set; ns, not significant; *, p<0.05; **, p<0.01; ***, p<0.001; ***, p<0.0001.

Upon comparison of shapes at the EC_100_ of each drug compared to the no-drug-treated control, we observed a significant increase in bacterial area and significant decrease in eccentricity in cells treated with D-cycloserine, lovastatin and pitavastatin (Figure 5C). For tetracycline, we found no significant differences in area or eccentricity compared to the untreated control group, likely because tetracycline does not directly interfere with cell wall biosynthesis. Thus, growth inhibition by statins causes bacterial shape defects that are similar to those caused by D-cycloserine, suggesting a shared mechanisms of bacterial growth inhibition resulting from targeting host isoprenoid and bacterial PG synthesis pathways.

## Discussion

Obligate intracellular pathogens such as *Rickettsia* are dependent upon nutrients and metabolites from their host cells as a consequence of reductive genome evolution (5, 6, 14). This suggests the possibility that this metabolic dependency represents an Achilles’ heel in the host-pathogen relationship, and that these pathogens could be sensitive to changes in host cell nutrient and metabolite levels. If metabolite availability is limited by the host or by extrinsic pressures such as chemical inhibition of a host biosynthetic pathway, pathogen growth should also be reduced. Here, we report the reduction of *R. parkeri* replication when the host isoprenoid pathway is inhibited with statins. This growth inhibition correlates with changes in bacterial shape that are consistent with defects in cell wall biosynthesis. Our results suggest that statins interfere with *Rickettsia* scavenging of host isoprenoids used for bacterial cell wall biosynthesis.

We observe that the isoprenoid biosynthesis pathway is under evolutionary flux in the order Rickettsiales, with the *Rickettsia* and *Orientia* lineages having lost genes encoding the upstream components of the MEP pathway, and *Rickettsia* having gained the *idi* gene. This is consistent with previous findings (14). Through our understanding of reductive genome evolution and gene acquisition in prokarykotes (5, 6, 8, 30), we can surmise a most parsimonious order of loss and gain of isoprenoid biosynthetic pathway genes in the Rickettsiales. We propose that the common ancestor of the *Rickettsia* and *Orientia* lineages, which inhabit the host cell cytoplasm, may have gained the ability to scavenge IPP and/or DMAPP from the cytoplasm by acquisition of a gene encoding transporter that moves these highly charged pyrophosphate molecules into the bacterial cell. There is evidence for the presence of such transporters in plants, bacteria, and protozoan parasites, although their molecular identities remain uncertain (31-33). The gain of a transporter would have enabled the loss of the upstream MEP pathway genes to streamline the bacterial genome and reduce fitness costs of pathway redundancy. Subsequently, the *idi* gene was gained in the *Rickettsia* lineage, which may reflect a need for additional DMAPP beyond that acquired from the host MEV pathway. In contrast, *Ehrlichia, Anaplasma, Wolbachia,* and *Neorickettsia* lineages, which grow within a membrane-bound vacuole, may not have sufficient access to host IPP and/or DMAPP and therefore have retained the entire MEP pathway.

We also found evidence of metabolic parasitism of host isoprenoids by *R. parkeri*. In particular, infected host cells become depleted of isoprenoid-derived storage forms of cholesterol, total cholesteryl esters and choleteryl oleate, perhaps to balance free cholesterol levels. Whether *Rickettsia* also scavenge downstream host isoprenoid products such as cholesterol, as observed for other bacteria such as *M. tuberculosis* and *Chlamydia trachomatis* (34-36), remains to be determined. Furthermore, *R. parkeri* produces downstream isoprenoid products even though it lacks genes for the upstream components of the MEP pathway. This metabolic parasitism suggests that inhibition of host isoprenoid biosynthesis should reduce the ability of the bacteria to synthesize bacterial isoprenoid products.

In keeping with this prediction, we found that statins, which inhibit host HMG-CoA-reductase, halt *R. parkeri* growth. Previous work had established an ability of statins to limit rickettsial plaque size (20), but mechanism of inhibition was not explored in detail. It was suggested that statins might inhibit *Rickettsia* adherence to and/or invasion of host cells based on previous studies that had found a role for cholesterol these processes (37, 38). Our work, however, reveals an effect of statins on intracellular bacterial growth, downstream of adherence/invasion. Furthermore, with increasing statin concentration, we find morphological changes in the shape and size of bacteria that mirror those caused by the cell wall synthesis inhibitor D-cycloserine. These observations lead us to conclude that statins inhibit bacterial growth by reducing host metabolite availability for production of bacterial products required for PG biosynthesis, leading to lethal defects in the bacterial cell wall.

Whether statins can be effective at preventing or treating human *Rickettsia* infections remains to be determined. Although statins are well tolerated in humans, and both statins tested in this study are FDA-approved, we have limited understanding of the dose response curves for inhibition of bacterial growth versus host toxicities that can include myopathy and increased risk of diabetes and stroke (39). Furthermore, it is unclear if statins have the same degree of bacterial growth inhibition in different rickettsial target cell types in vivo. Additional studies in animal models will shed light on future promise of statins as a prophylaxis or treatment for rickettsial infections. Currently, cases of rickettsial diseases are effectively treated with tetracyclines, although there are reported cases of tetracycline-resistant scrub typhus (40, 41), and tetracycline treatment is contraindicated for pregnant women and young children (42). Because a limited number of antibiotics are effective in treating rickettsial infections (22), host-targeted therapeutics would be useful additions to treatment regimens. Furthermore, targeting host biosynthesis of essential metabolites may limit the development of antibiotic resistance.

Beyond the scope of rickettsial infections, this study points to a potential to use host-targeted therapeutics for a wide variety of infectious microbes, including viruses, bacteria, fungi, and parasites. Similar to *R. parkeri*, the parasite *Cryptosporidium parvum* lacks the canonical protozoan isoprenoid pathway and has been shown to be sensitive to statins (43). In the future, bioinformatic analyses may identify other intracellular bacteria that have incomplete upstream isoprenoid pathways and intact downstream isoprenoid pathways, and thus may also be sensitive to statins. Furthermore, *Rickettsia* and other parasites may be sensitive to chemical inhibition of other host metabolic pathways. Layering our knowledge of FDA-approved compounds that target host biosynthetic pathways onto predicted metabolic pathways hijacked by pathogens may therefore enable the systematic identification of drug classes that could be re-purposed as antibiotics.

## Materials and Methods

### Reagents

Hoechst 33342 (B2261), D-cycloserine (C6880), tetracycline (T7660), fosmidomycin (F8682), mevalonate (M4667), and alendronate sodium hydrate (A4978) were obtained from Sigma-Aldrich. ProLong Gold Antifade (P36930), Pitavastatin (494210), Lovastatin InSolution (4381875), goat anti-mouse Alexa-488 secondary antibody (R37120), 0.25% Trypsin-EDTA phenol red (25200056) and Dynamo HS SYBR Green qPCR kit (F-410L) were obtained from Thermo-Fisher Scientific. Anti-*Rickettsia* 14-13 antibody was generously provided by Ted Hackstadt, NIH/NIAID Rocky Mountain Laboratories (44).

### Cell culture and R. parkeri infections

Confluent low-passage African green monkey epithelial Vero cells were obtained from the UC Berkeley Cell Culture Facility and grown at 37°C, 5% CO_2_, in high glucose (4.5 g/l) DMEM (Gibco, Life Technologies, 11965092) supplemented with 2% fetal bovine serum (FBS, Benchmark). *R. parkeri* strain Portsmouth was generously provided by Chris Paddock (Centers for Disease Control and Prevention). *R. parkeri* was propagated by infecting monolayers of Vero cells with an MOI = 0.1 of wild-type *R. parkeri* and growing them at 33°C, 5% CO_2_ in DMEM plus 2% FBS. Bacteria were purified from infected cells as described previously (45). Briefly, infected cells were lysed by dounce homogenization in cold K-36 buffer (0.05 M KH_2_PO_4_, 0.05 M K_2_HPO_4_, pH 7, 100 mM KCl, 15 mM NaCl) to release bacteria, the lysate was overlaid onto 30% MD-76R (Mallinckrodt Inc., 1317-07), centrifuged at 58,300 x g for 20 min at 4°C in a SW-28 swinging bucket rotor, and resuspended in cold brain heart infusion (BHI) media (BD Difco, 237500) broth. Aliquots of bacteria were immediately frozen at −80°C after purification and each infection was initiated from a single thawed aliquot of bacteria.

### Phylogenetic analyses, idi locus mapping, and whole genome alignments

Phylogenetic relationships between bacterial species were analyzed using 16S rDNA sequences downloaded from the SILVA database (https://www.arb-silva.de/). Phylogenetic trees were built with the Tree Builder program on Geneious software version 9.1.8 using the neighbor-joining method with *Pelagibacter ubique* as the outgroup. The presence/absence of the MEP pathway or *idi* gene was determined using the KEGG pathway database (http://www.kegg.jp/) and manual annotations of the terpenoid backbone biosynthesis pathway (http://www.kegg.jp/kegg-bin/show_pathway?map=map00900) for each bacterial species. Determination of intracellular niche was performed by manual literature curation. *Idi* locus map and annotations were examined using Geneious software version 9.1.8 using the *R. parkeri* str. Portsmouth genome sequence (NCBI RefSeq: NC_017044.1). Whole genome alignments of *R. parkeri* str. Portsmouth and *R. rickettsii* str. Iowa (NCBI RefSeq: NC_010263.3) were performed using the Mauve 2.3.1 (46) plugin in Geneious software using the progressive Mauve algorithm with automatically-calculated-minimum locally collinear block (LCBs) scores.

### Mass spectrometry

Monolayers of Vero cells plated in 6-well plates were infected (or an uninfected control) with *R. parkeri* as described above and were incubated for 4 d at a 33°C 5% CO_2_. On the day of harvest, cells in each well were washed with 4 ml of 1 x phosphate buffered saline (PBS). 1 ml of fresh 1 x PBS was added to each well and used to scrape the cells, centrifuged at 1000 x g for 5 min on a microfuge and the supernatant was aspirated from the cell pellet. Cell pellets were extracted in 2:1:1 chloroform:methanol:PBS solution with addition of 10 nM dodecylglycerol internal standard, after which the organic phase was separated, collected, and dried down under N_2_ gas. The dried-down lipidome extract was then resuspended in 150 µl chloroform and stored at −80°C until analysis. For mass spectrometry analysis, an aliquot of this sample solution was injected into an Agilent 6430 LC-MS/MS and metabolites were chromatographically separated and analyzed by SRM-based targeted LC-MS/MS as previously described (47, 48).

### Immunofluorescence microscopy

To measure bacterial shape, Vero cells were plated at 50% confluency on 96-well glass bottom plates in DMEM plus 2% FBS and allowed to settle overnight at 37°C, 5% CO_2_. The following day, cells were infected with an MOI = 0.1 of wild-type *R. parkeri*, centrifuged at 300 x *g* for 5 min and incubated for 2 h at 33°C, 5% CO_2_. Media containing bacteria was aspirated and replaced with media with or without the appropriate concentration of drug (see above), and cells were further incubated for 72 h at 33°C, 5% CO_2_. Cells were fixed with 4% paraformaldehyde in 1 x PBS for 20 min at room temperature, washed with 1 x PBS, and permeabilized with 0.05% Triton X-100 in 1 x PBS for 5 min. Cells were then stained for immunofluorescence with mouse anti-*Rickettsia* primary antibody 14-13 and goat anti-mouse Alexa-488 secondary antibody to stain bacteria, and Hoechst 33342 to stain DNA. Infected cells were imaged on a Nikon Ti Eclipse microscope with a Yokogawa CSU-XI spinning disc confocal, 100X (1.4 NA) Plan Apo objective, a Clara Interline CCD Camera and MetaMorph software taking 0.15 µm z-slices across 5 µm in the z-plane. Maximum Z-projections for each channel were made using ImageJ and a custom pipeline was created in CellProfiler software (29) to identify individual bacteria in each image. The CellProfiler module MeasureObjectSizeShape was used to calculate size and shape parameters of each bacterium.

### R. parkeri qPCR growth and drug dose response curves

Confluent Vero cells grown in 96-well tissue culture plates were infected with an MOI = 0.1 of *R. parkeri* and incubated in DMEM plus 2% FBS with or without the appropriate concentration of drug (see above) for 72 h at 33°C, 5% CO_2_. To harvest cells at the appropriate time point, media was aspirated and cells lifted with 50 µl of 0.25% trypsin-EDTA followed by incubation at 37°C for 5 min. Lifted cells were resuspended with an additional 50 µl of DMEM before adding to 50 µl Nuclei Lysis Buffer (Wizard Genomic DNA Purification kit; Promega) and frozen at −20°C overnight. Cells were then thawed and boiled for 10 min to release genomic DNA. To remove RNA, 20 µg/mL RNAse A was added to each sample, incubated for 15 min at 37°C, and then samples were cooled to room temperature. Protein was removed by adding 50 µl of protein precipitation solution, mixing by pipetting, then centrifuging in a microfuge for 15 min at 1500 x *g* at 4°C. DNA was precipitated by adding 100 µl of the resulting supernatant to 100 µl of isopropanol, mixing by pipetting, and centrifuging for 15 min at 1500 x *g* at 4°C. Isopropanol was removed and the DNA pellet was washed with 70% ethanol and centrifuged for 15 min at 1500 x *g*. Resulting DNA pellets were dried, then resuspended in 50 µl of H_2_O and allowed to rehydrate overnight at 4°C. For quantitative real-time PCR, 5 µl genomic DNA was used with primers to the *R. parkeri* gene encoding the 17kDa antigen (49) on a Bio-Rad CFX96 Touch Real-Time PCR Detection System. *Rickettsia* genome copy number was quantified against a standard curve of a plasmid containing a single copy of the *R. parkeri* 17kDa gene. Regression analysis and EC50 calculations were performed using GraphPad Prism 7 software.

## Statistical analyses

Statistical analysis was performed in GraphPad PRISM 7 and statistical parameters and significance are reported in the Figures and Figure Legends. Statistical significance was determined either by an unpaired Student’s t-test or a one-way ANOVA where indicated. Statistical significance was denoted by asterisks as follows: *, p < 0.05; **, p < 0.01; ***, p < 0.001; ****, p < 0.0001, compared to controls.

## Acknowledgments

We are grateful to Rebecca Lamason for critical reading of the manuscript. We thank Eric Lee and Dan Portnoy for reagents. We thank Matthew Knight, Eric Lee, Ellen Yeh, David Booth, Thibaut Brunet, and members of the Welch lab for helpful technical advice and discussions. This work was performed in part at the CRL MIC, which is supported by NIH grant S10RR027696-01. V.A. was supported by the NIH/NIAID grant F32 AI133912A. M.D.W. was supported by NIH/NIAID grants R01 AI109044 and R21 AI109270. C.A.B. and D.K.N. were supported by NIH/NCI grant R01 CA172667.

## References

1. Yu X-J, Walker DH. 2006. The Order Rickettsiales, p 493–528. In The Prokaryotes. Springer New York, New York, NY.

2. Kirkland KB, Wilkinson WE, Sexton DJ. 1995. Therapeutic delay and mortality in cases of Rocky Mountain spotted fever. Clin Infect Dis 20:1118–1121.

3. Paddock CD, Sumner JW, Comer JA, Zaki SR, Goldsmith CS, Goddard J, McLellan SLF, Tamminga CL, Ohl CA. 2004. *Rickettsia parkeri*: a newly recognized cause of spotted fever rickettsiosis in the United States. Clin Infect Dis 38:805–811.

4. Paddock CD, Finley RW, Wright CS, Robinson HN, Schrodt BJ, Lane CC, Ekenna O, Blass MA, Tamminga CL, Ohl CA, McLellan SLF, Goddard J, Holman RC, Openshaw JJ, Sumner JW, Zaki SR, Eremeeva ME. 2008. *Rickettsia parkeri* rickettsiosis and its clinical distinction from Rocky Mountain spotted fever. Clin Infect Dis 47:1188–1196.

5. Andersson JO, Andersson SG. 1999. Insights into the evolutionary process of genome degradation. Curr Opin Genet Dev 9:664–671.

6. Moran NA. 2002. Microbial minimalism: genome reduction in bacterial pathogens. Cell 108:583–586.

7. Darby AC, Cho N-H, Fuxelius H-H, Westberg J, Andersson SGE. 2007. Intracellular pathogens go extreme: genome evolution in the Rickettsiales. Trends Genet 23:511–520.

8. Blanc G, Ogata H, Robert C, Audic S, Suhre K, Vestris G, Claverie J-M, Raoult D. 2007. Reductive genome evolution from the mother of *Rickettsia*. PLoS Genet 3:e14.

9. Gillespie JJ, Williams K, Shukla M, Snyder EE, Nordberg EK, Ceraul SM, Dharmanolla C, Rainey D, Soneja J, Shallom JM, Vishnubhat ND, Wattam R, Purkayastha A, Czar M, Crasta O, Setubal JC, Azad AF, Sobral BS. 2008. *Rickettsia* phylogenomics: unwinding the intricacies of obligate intracellular life. PLoS ONE 3:e2018.

10. Heuston S, Begley M, Gahan CGM, Hill C. 2012. Isoprenoid biosynthesis in bacterial pathogens. Microbiology 158:1389–1401.

11. Goldstein JL, Brown MS. 1990. Regulation of the mevalonate pathway. Nature 343:425–430.

12. Eberl M, Hintz M, Jamba Z, Beck E, Jomaa H, Christiansen G. 2004. *Mycoplasma penetrans* is capable of activating V gamma 9/V delta 2 T cells while other human pathogenic mycoplasmas fail to do so. Infect Immun 72:4881–4883.

13. Clastre M, Goubard A, Prel A, Mincheva Z, Viaud-Massuart M-C, Bout D, Rideau M, Velge-Roussel F, Laurent F. 2007. The methylerythritol phosphate pathway for isoprenoid biosynthesis in coccidia: presence and sensitivity to fosmidomycin. Exp Parasitol 116:375–384.

14. Driscoll TP, Verhoeve VI, Guillotte ML, Lehman SS, Rennoll SA, Beier-Sexton M, Rahman MS, Azad AF, Gillespie JJ. 2017. Wholly *Rickettsia*! Reconstructed metabolic profile of the quintessential bacterial parasite of eukaryotic cells. MBio 8:84.

15. Berthelot K, Estevez Y, Deffieux A, Peruch F. 2012. Isopentenyl diphosphate isomerase: A checkpoint to isoprenoid biosynthesis. Biochimie 94:1621–1634.

16. Murphy DJ. 2001. The biogenesis and functions of lipid bodies in animals, plants and microorganisms. Prog Lipid Res 40:325–438.

17. Manat G, Roure S, Auger R, Bouhss A, Barreteau H, Mengin-Lecreulx D, Touzé T. 2014. Deciphering the metabolism of undecaprenyl-phosphate: the bacterial cell-wall unit carrier at the membrane frontier. Microb Drug Resist 20:199–214.

18. Nowicka B, Kruk J. 2010. Occurrence, biosynthesis and function of isoprenoid quinones. Biochim Biophys Acta 1797:1587–1605.

19. Aussel L, Pierrel F, Loiseau L, Lombard M, Fontecave M, Barras F. 2014. Biosynthesis and physiology of coenzyme Q in bacteria. Biochim Biophys Acta 1837:1004–1011.

20. Botelho-Nevers E, Rolain JM, Espinosa L, Raoult D. 2008. Statins limit *Rickettsia conorii* infection in cells. Int J Antimicrob Agents 32:344–348.

21. Reed SCO, Serio AW, Welch MD. 2012. *Rickettsia parkeri* invasion of diverse host cells involves an Arp2/3 complex, WAVE complex and Rho-family GTPasedependent pathway. Cell Microbiol 14:529–545.

22. Rolain JM, Maurin M, Vestris G, Raoult D. 1998. In vitro susceptibilities of 27 rickettsiae to 13 antimicrobials. Antimicrob Agents Chemother 42:1537–1541.

23. Pang H, Winkler HH. 1994. Analysis of the peptidoglycan of *Rickettsia prowazekii*. J Bacteriol 176:923–926.

24. Baisa G, Stabo NJ, Welch RA. 2013. Characterization of *Escherichia coli* D- cycloserine transport and resistant mutants. J Bacteriol 195:1389–1399.

25. Desjardins CA, Cohen KA, Munsamy V, Abeel T, Maharaj K, Walker BJ, Shea TP, Almeida DV, Manson AL, Salazar A, Padayatchi N, O’Donnell MR, Mlisana KP, Wortman J, Birren BW, Grosset J, Earl AM, Pym AS. 2016. Genomic and functional analyses of *Mycobacterium tuberculosis* strains implicate *ald* in D-cycloserine resistance. Nat Genet 48:544–551.

26. Delcour AH. 2009. Outer membrane permeability and antibiotic resistance. Biochim Biophys Acta 1794:808–816.

27. Wisseman CL, Silverman DJ, Waddell A, Brown DT. 1982. Penicillin-induced unstable intracellular formation of spheroplasts by rickettsiae. J Infect Dis 146:147–158.

28. Allan EJ, Hoischen C, Gumpert J. 2009. Bacterial L-forms. Adv Appl Microbiol 68:1–39.

29. Carpenter AE, Jones TR, Lamprecht MR, Clarke C, Kang IH, Friman O, Guertin DA, Chang JH, Lindquist RA, Moffat J, Golland P, Sabatini DM. 2006. CellProfiler: image analysis software for identifying and quantifying cell phenotypes. Genome Biol 7:R100.

30. Wolf YI, Koonin EV. 2013. Genome reduction as the dominant mode of evolution. Bioessays 35:829–837.

31. Bick JA, Lange BM. 2003. Metabolic cross talk between cytosolic and plastidial pathways of isoprenoid biosynthesis: unidirectional transport of intermediates across the chloroplast envelope membrane. Arch Biochem Biophys 415:146–154.

32. Flügge U-I, Gao W. 2005. Transport of isoprenoid intermediates across chloroplast envelope membranes. Plant Biol 7:91–97.

33. Yeh E, DeRisi JL. 2011. Chemical rescue of malaria parasites lacking an apicoplast defines organelle function in blood-stage *Plasmodium falciparum*. PLoS Biol 9:e1001138.

34. Carabeo RA, Mead DJ, Hackstadt T. 2003. Golgi-dependent transport of cholesterol to the *Chlamydia trachomatis* inclusion. Proc Natl Acad Sci U S A 100:6771–6776.

35. Samanta D, Mulye M, Clemente TM, Justis AV, Gilk SD. 2017. Manipulation of host cholesterol by obligate intracellular bacteria. Front Cell Infect Microbiol 7:165.

36. Wilburn KM, Fieweger RA, VanderVen BC. 2018. Cholesterol and fatty acids grease the wheels of *Mycobacterium tuberculosis* pathogenesis. Pathog Dis 76.

37. Ramm LE, Winkler HH. 1976. Identification of cholesterol in the receptor site for rickettsiae on sheep erythrocyte membranes. Infect Immun 13:120–126.

38. Martinez JJ, Seveau S, Veiga E, Matsuyama S, Cossart P. 2005. Ku70, a component of DNA-dependent protein kinase, is a mammalian receptor for *Rickettsia conorii*. Cell 123:1013–1023.

39. Bellosta S, Corsini A. 2018. Statin drug interactions and related adverse reactions: an update. Expert Opin Drug Saf 17:25–37.

40. Strickman D, Sheer T, Salata K, Hershey J, Dasch G, Kelly D, Kuschner R. 1995. In vitro effectiveness of azithromycin against doxycycline-resistant and - susceptible strains of *Rickettsia tsutsugamushi*, etiologic agent of scrub typhus. Antimicrob Agents Chemother 39:2406–2410.

41. Kelly DJ, Fuerst PA, Richards AL. 2017. The historical case for and the future study of antibiotic-resistant scrub typhus. Trop Med Infect Dis 2:63.

42. Cross R, Ling C, Day NPJ, McGready R, Paris DH. 2016. Revisiting doxycycline in pregnancy and early childhood--time to rebuild its reputation? Expert Opin Drug Saf, 29 ed. 15:367–382.

43. Bessoff K, Sateriale A, Lee KK, Huston CD. 2013. Drug repurposing screen reveals FDA-approved inhibitors of human HMG-CoA reductase and isoprenoid synthesis that block *Cryptosporidium parvum* growth. Antimicrob Agents Chemother 57:1804–1814.

44. Anacker RL, Mann RE, Gonzales C. 1987. Reactivity of monoclonal antibodies to *Rickettsia rickettsii* with spotted fever and typhus group rickettsiae. J Clin Microbiol 25:167–171.

45. Lamason RL, Bastounis E, Kafai NM, Serrano R, Del Álamo JC, Theriot JA, Welch MD. 2016. Rickettsia Sca4 reduces vinculin-mediated intercellular tension to promote spread. Cell 167:670–683.e10.

46. Darling AE, Mau B, Perna NT. 2010. progressiveMauve: multiple genome alignment with gene gain, loss and rearrangement. PLoS ONE 5:e11147.

47. Benjamin DI, Cozzo A, Ji X, Roberts LS, Louie SM, Mulvihill MM, Luo K, Nomura DK. 2013. Ether lipid generating enzyme AGPS alters the balance of structural and signaling lipids to fuel cancer pathogenicity. Proc Natl Acad Sci USA 110:14912–14917.

48. Louie SM, Grossman EA, Crawford LA, Ding L, Camarda R, Huffman TR, Miyamoto DK, Goga A, Weerapana E, Nomura DK. 2016. GSTP1 is a driver of triple-negative breast cancer cell metabolism and pathogenicity. Cell Chem Biol 23:567–578.

49. Grasperge BJ, Reif KE, Morgan TD, Sunyakumthorn P, Bynog J, Paddock CD, Macaluso KR. 2012. Susceptibility of inbred mice to *Rickettsia parkeri*. Infect Immun 80:1846–1852.

