## Supplemental Table 1 for "A metabolic dependency for host isoprenoids in the obligate intracellular pathogen *Rickettsia parkeri* underlies a sensitivity for the statin class of host-targeted therapeutics"

**Supplemental Table S1**

| Locus tag | Product | Position | Length | Protein ID |
|---|---|---|---|---|
| MC1_RS03625 | hypothetical protein | 694497 | 1712 | pseudogene |
| MC1_RS03630 | DNA methyltransferase | 696210 | 813 | pseudogene |
| MC1_RS03635 | sensor histidine kinase | 697402 | 912 | WP_014410747.1 |
| MC1_RS03640 | bifunctional (p)ppGpp synthetase/guanosine-3',5'-bis(diphosphate) 3'- | 698450 | 681 | WP_014410748.1 |
| MC1_RS03645 | hypothetical protein | 699600 | 2871 | WP_014410749.1 |
| MC1_RS03650 | preprotein translocase subunit SecA | 702483 | 6672 | WP_014410750.1 |
| MC1_RS03655 | conjugal transfer protein TraD | 709338 | 276 | WP_041472287.1 |
| MC1_RS03660 | IS256 family transposase | 709980 | 1216 | pseudogene |
| MC1_RS07740 | hypothetical protein | 711323 | 646 | pseudogene |
| MC1_RS07745 | hypothetical protein | 712056 | 219 | pseudogene |
| MC1_RS07035 | reverse transcriptase | 712487 | 573 | WP_014410755.1 |
| MC1_RS03670 | conjugal transfer protein TraA | 713785 | 4137 | WP_014410756.1 |
| MC1_RS03675 | transposase | 717969 | 957 | WP_041472215.1 |
| MC1_RS03680 | conjugal transfer protein TraD | 719163 | 777 | WP_081497779.1 |
| MC1_RS07750 | conjugal transfer protein TraD | 719946 | 276 | pseudogene |
| MC1_RS03685 | hypothetical protein | 720422 | 399 | WP_014410759.1 |
| MC1_RS03690 | transposase | 720858 | 1044 | WP_014410760.1 |
| MC1_RS03695 | DUF87 domain-containing protein | 722084 | 1695 | WP_014410761.1 |
| MC1_RS03700 | hypothetical protein | 723783 | 1195 | pseudogene |
| MC1_RS07755 | hypothetical protein | 724964 | 408 | WP_014410762.1 |
| MC1_RS03705 | F pilus assembly protein TraH | 725338 | 894 | WP_014410763.1 |
| MC1_RS03710 | conjugal transfer protein TraF | 726221 | 462 | pseudogene |

**Supplemental Table S1:** Genes missing in *R. rickettsii* adjacent to the *R. parkeri idi* locus.
List of 22 genes, annotation, sequence start site, length, and protein ID missing in the R. rickettsii genome as discovered by Mauve whole genome alignment to R. parkeri.
